# Targeting deubiquitinating enzymes (DUBs) and ubiquitin pathway modulators to enhance host defense against bacterial infections

**DOI:** 10.1101/2025.01.27.635188

**Authors:** John Santelices, Alexander Schultz, Alyssa Walker, Nicole Adams, Deyaneira Tirado, Hailey Barker, Aria Eshraghi, Daniel M. Czyż, Mariola J. Ferraro

**Author notes:** Correspondence: Mariola J. Ferraro, Microbiology and Cell Science Department, University of Florida, Gainesville, FL 32611, USA.

## Abstract

The rise of antibiotic-resistant bacterial pathogens poses a critical global health challenge, necessitating innovative therapeutic approaches. This study explores host-targeted therapies (HTTs) by focusing on deubiquitinating enzymes (DUBs), essential modulators of the ubiquitin-proteasome system (UPS) that regulate host-pathogen interactions during many bacterial infections. Using *Salmonella*-infected macrophages as a model, we identified UPS modulators that enhance bacterial clearance and observed significant changes in DUB expression, particularly USP25, USP46, and Otud7b. The small-molecule DUB inhibitor AZ-1 significantly reduced intracellular bacterial loads in vitro and mitigated early disease severity in a murine model by decreasing fecal bacterial loads and preserving host weight. However, AZ-1 alone did not achieve complete clearance of *Salmonella* and required combination with extracellular-targeting antibiotics for optimal efficacy. Notably, AZ-1 demonstrated broad-spectrum activity against multidrug-resistant pathogens, including *Pseudomonas aeruginosa, Klebsiella pneumoniae*, and *Acinetobacter baumannii.* Transcriptomic analyses revealed infection-induced DUB regulation and highlighted pathways modulating immune responses, including TNF-α secretion. These findings highlight the potential of targeting the UPS as a host-directed antimicrobial strategy and provide a foundation for developing innovative therapies to combat antimicrobial resistance.

## 1. Introduction

Infectious diseases continue to pose a significant global health threat, with bacterial pathogens constantly evolving new strategies to evade host immune defenses. Among these pathogens, *Salmonella* spp. are particularly concerning due to their ability to cause severe, often life-threatening infections. This issue is compounded by the increasing prevalence of antimicrobial resistance (AMR), which renders conventional treatment ineffective (1, 2). The rising AMR in Gram-negative bacteria like *Salmonella* fuels the urgent need for novel therapeutic approaches that extend beyond traditional antibiotics. One promising approach to combat resistant pathogens involves targeting components of the host’s cellular machinery, such as the ubiquitin-proteasome system (UPS). The UPS is crucial for maintaining protein homeostasis, DNA repair, and cell cycle regulation, which are essential for maintaining cellular function (3). Pathogens have evolved sophisticated strategies to hijack this system, particularly through manipulation of deubiquitinating enzymes (deubiquitinases; DUBs) (4).

*Salmonella*, an intracellular pathogen, engages unique mechanisms to survive in various cell types, including M cells, epithelial cells, macrophages, and dendritic cells (DCs) [reviewed in (5)]. Although macrophages are central to the host’s innate immune defense, they paradoxically serve as a Trojan horse for *Salmonella* and support its replication (6–9). This phenomenon is facilitated by the bacterium’s ability to reprogram host cells such that it can survive within a specialized vacuole (10–12). The intracellular survival of *Salmonella* depends on effector proteins encoded from *Salmonella* pathogenicity islands 1 and 2 (SPI-1 and SPI-2), which manipulate host cell processes, immune responses, and cell signaling pathways, including those regulated by the UPS [reviewed in (13, 14)]. Treating *Salmonella* infections is becoming progressively more challenging due to its rising AMR,(1, 2) making it imperative to explore alternative therapeutic strategies. Host-targeted therapies (HTTs) offer a promising approach to control infection by disrupting the cellular pathways that pathogens rely on for survival, thereby reducing the selective pressure for antibiotic resistance (15, 16). This approach offers a potential solution to address the escalating AMR crisis (16). Given the central role of the UPS in regulating host-pathogen interactions, targeting specific components of this system—particularly DUBs—could effectively impair the ability of *Salmonella* to survive and replicate within host cells(17). This study evaluates the therapeutic potential of DUB inhibitors and other UPS modulators as a host-targeting approach to clearing bacteria that could bring innovative treatments against *Salmonella* and other bacterial pathogens (18).

DUBs, central to the UPS, are responsible for removing ubiquitin and ubiquitin-like modifiers from substrates, reversing post-translational modifications, modulating signaling pathways, and replenishing the pool of free ubiquitin necessary for ongoing ubiquitination processes (19). This diverse functionality impacts numerous cellular functions, including protein degradation, cellular localization, and protein-protein interactions. Through these mechanisms, DUBs influence a wide array of biological processes, including those related to human health, viral infections, and pathogen-host interactions (20–24). Notably, pathogenic bacteria, such as the intracellular pathogen *Salmonella*, exploit the host’s ubiquitination machinery to evade immune responses and establish persistent infections (18, 25). Targeting DUBs within the ubiquitin system presents a possible avenue for therapeutic development, potentially disrupting the ability of bacteria to survive and replicate within host cells (18).

In this study, we explored the therapeutic potential of UPS-specific inhibitors as a host-targeted strategy for bacterial infections. We identified key molecular targets and promising chemical inhibitors capable of combating pathogens such as *Salmonella* and ESKAPE pathogens, including such pathogens as *Klebsiella pneumoniae, Acinetobacter baumannii*, or *Pseudomonas aeruginosa*. To validate the therapeutic potential of DUB inhibitors, we performed a proof-of-principle study using a murine infection model. Our results demonstrated that AZ-1 treatment conferred moderate protective effects by reducing bacterial colonization and weight loss as well as improving body scores during *Salmonella* infection. While AZ-1 alone was insufficient to prevent infection-related lethality in animals, its efficacy could be potentially enhanced with antibiotics targeting extracellular bacteria. In addition to these findings, we conducted a screen of commercially available UPS modulators and DUB inhibitors from a curated library, identifying their roles in modulating host immune responses and restricting intracellular bacterial replication. These results support the potential of targeting the UPS to enhance host immunity towards more effective combination therapeutic strategies against bacterial infections, addressing both intracellular and extracellular pathogens.

## 2. Results

### 2.1 Small molecule modulators targeting the ubiquitin pathway as potential compounds enhancing macrophage-mediated bacterial clearance

We explored the impact of inhibiting specific components of the UPS on *Salmonella* pathogenicity using a high-throughput screening (HTS) assay. Using this assay, we screened a library of 257 small molecule modulators targeting key enzymes in the ubiquitin pathway, aiming to identify compounds that enhance macrophage-mediated bacterial clearance without affecting cell viability. We also assessed whether these compounds directly affected *Salmonella* viability in axenic growth assays and evaluated their impact on pro-inflammatory cytokine TNF-α levels during infection.

In the initial assay, macrophages infected with GFP-labeled *Salmonella* Typhimurium UK-1 were seeded into 96-well plates. Hoechst stain was used to label host cell nuclei, while HCS CellMask Red highlighted the cytoplasm, enabling precise quantification of intracellular bacteria while minimizing interference from extracellular bacteria (**Fig. 1A**). Following infection, the cells were treated with each of the 257 UPS-targeting compounds, and the macrophages were incubated to allow bacterial uptake. High-content imaging was used to capture and quantify the number of intracellular bacteria per cell, assessing the efficacy of each compound. The HTS assay revealed 59 compounds that significantly reduced intracellular bacterial counts compared to untreated controls (**Fig. 1B**, **Table 1**), with some compounds reaching over 10-fold reduction and showing high statistical significance.

**Figure 1.**
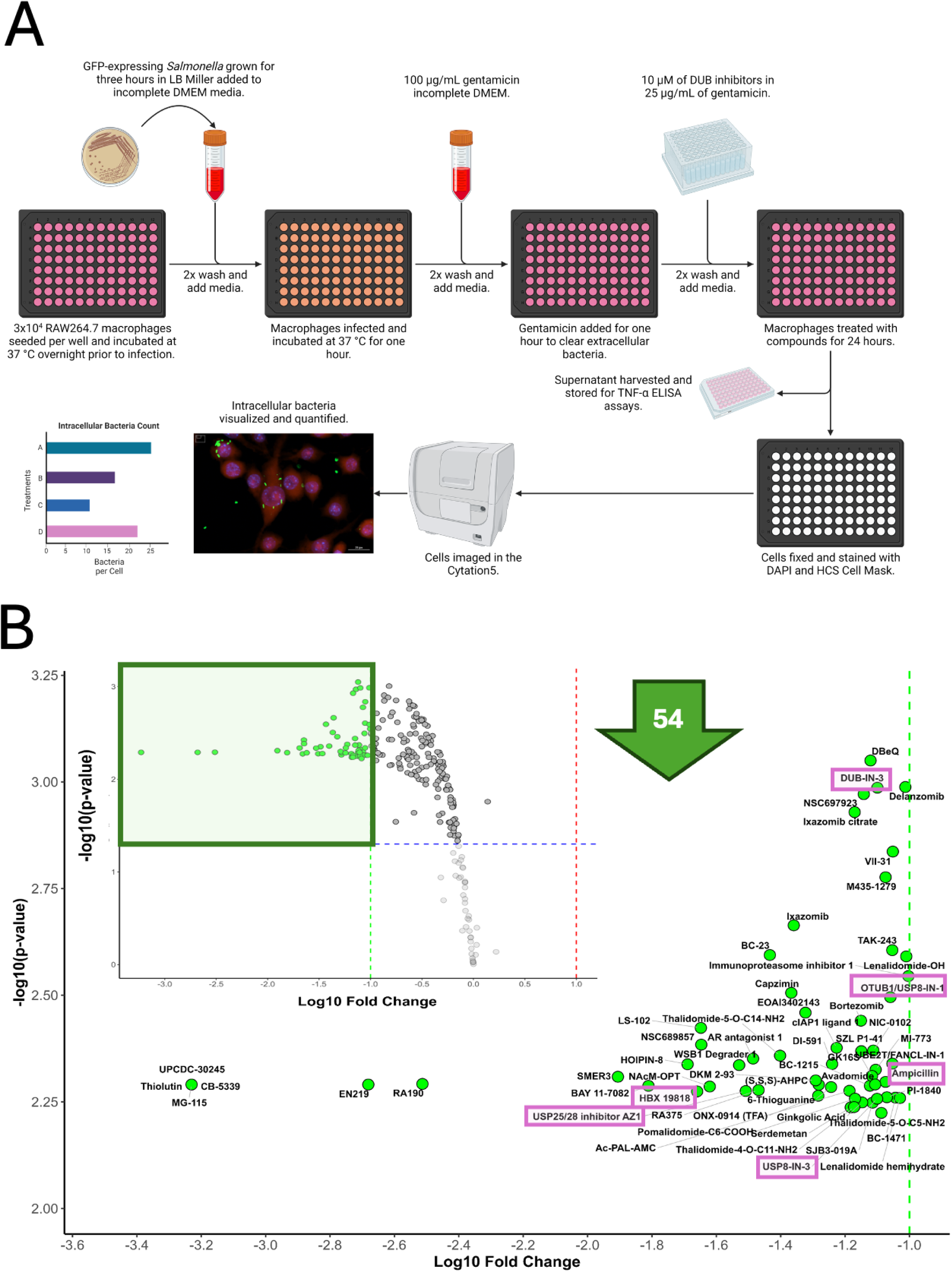
Screening of modulators of the Ubiquitin Proteasome System (UPS) pathway for compounds supporting macrophage-mediated bacterial clearance. **(A)** Schematic representation of the high-throughput assay designed to identify compounds that enhance macrophage-mediated clearance of *Salmonella* Typhimurium strain UK-1 infection (MOI 30:1). The Ubiquitination Compound Library (MedChem Express, *HY-L050*) was used for screening at a 10 µM concentration. This library comprises 257 small molecule modulators targeting key enzymes in the ubiquitin pathway. **(B)** The impact of UPS pathway inhibitors on intracellular *Salmonella* was assessed. Intracellular bacterial counts were measured at 24 hours post-infection (hpi) with *Salmonella* (MOI 30:1) using a high-throughput screening (HTS) assay and normalized to cell count. Data were analyzed in R, with mean replicate values calculated for each treatment. Fold changes and p-values compared to vehicle controls were determined using t-tests, followed by Benjamini-Hochberg correction for multiple comparisons. Significant modulators of *Salmonella* infection were visualized using a volcano plot, indicating the log10 fold change and -log10(p-value).

**Table 1.**
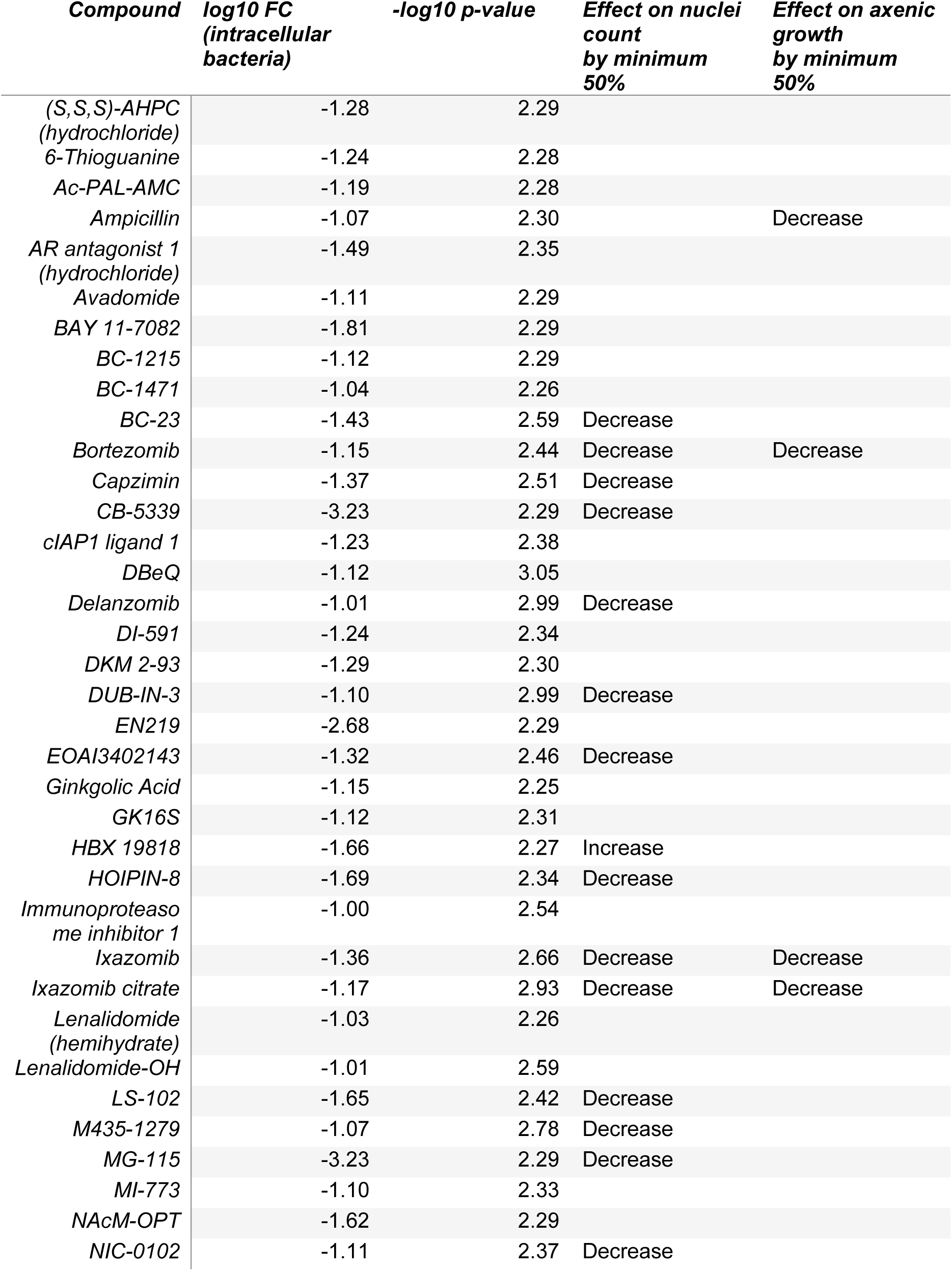
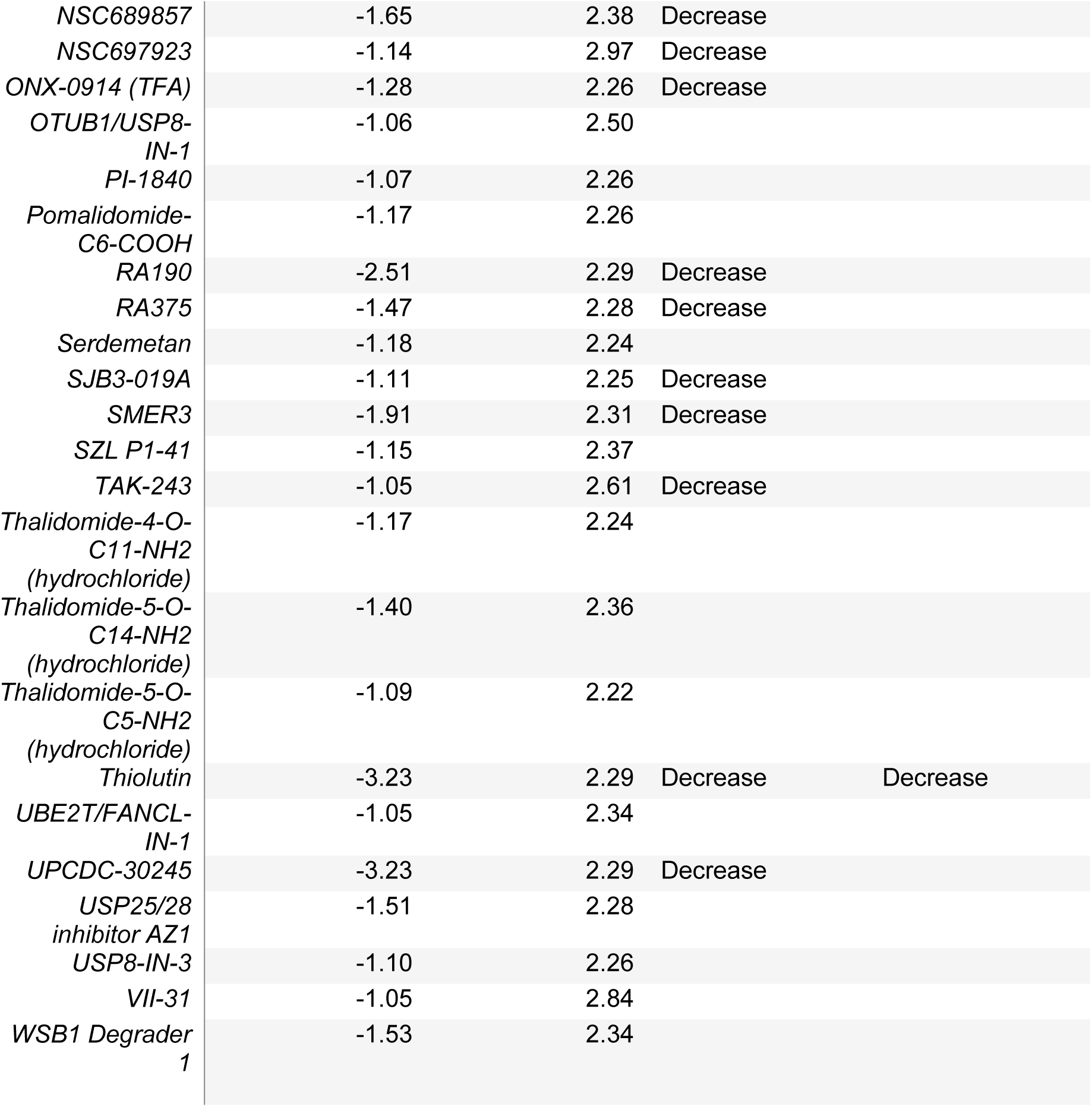
Effects of ubiquitin pathway-modulating compounds on macrophage-mediated bacterial clearance, cell number, and axenic growth. Compounds that enhance bacterial clearance without inducing significant macrophage cytotoxicity are highlighted in bold for their therapeutic potential.

Further analysis revealed that some compounds that decreased intramacrophage survival of *Salmonella* also adversely affected host viability, therefore not being feasible host-targeting targets. These compounds included Bortezomib, Capzimin, Ixazomib citrate, Delanzomib, EOAI3402143, MG-115, NIC-0102, NSC689857, and NSC697923, as indicated by a decrease in nuclei count by at least 50% in presence of *Salmonella*. In contrast, HBX 19818 maintained host cell viability—showing no decrease or even a slight improvement compared to infected, untreated cells—while simultaneously reducing the bacterial load (**Fig. 2A**).

**Figure 2.**
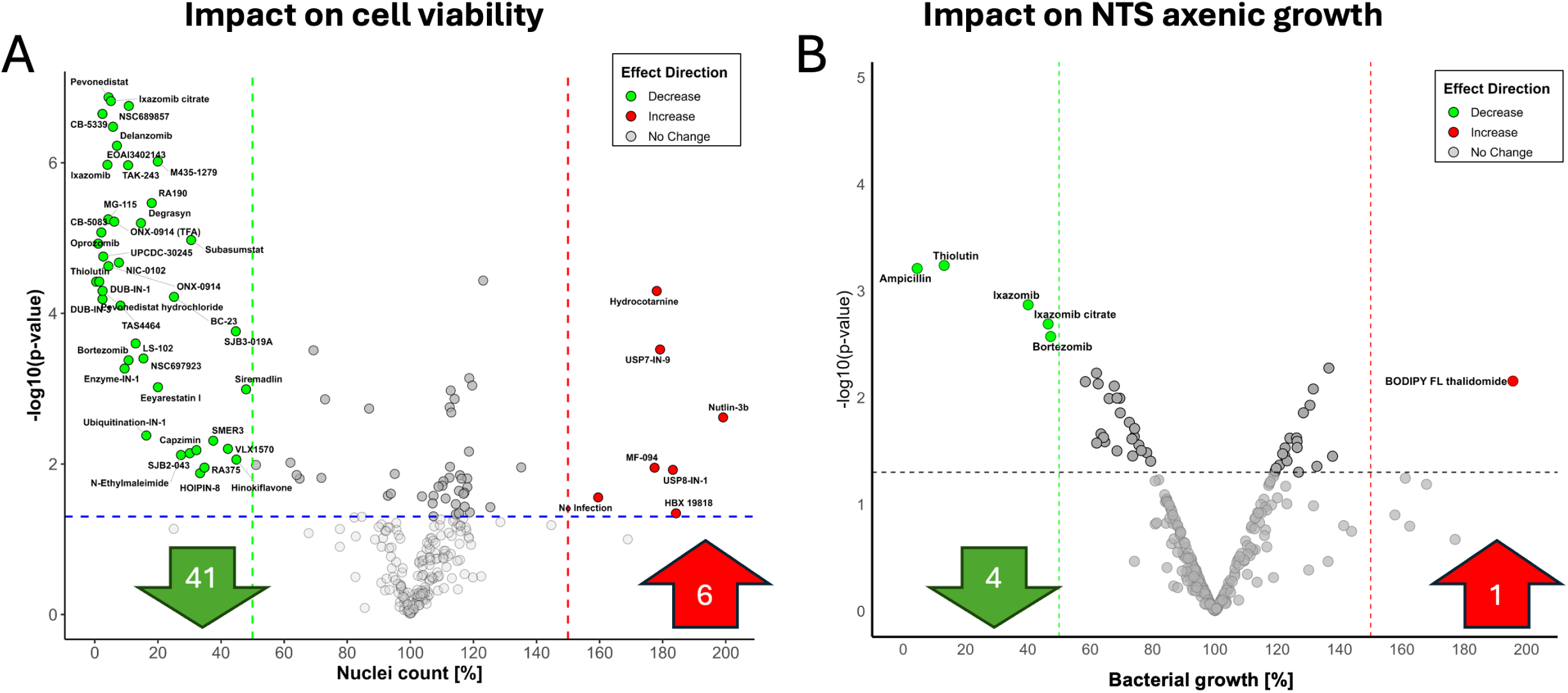
Screening of Ubiquitin Proteasome System (UPS) pathway modulators for compounds that decrease cell count and inhibit bacterial growth in axenic culture. **(A)** An experiment was performed as in **Fig 1B**. Nuclei count of macrophages treated with UPS pathway inhibitors at a concentration of 10 µM. The Ubiquitination Compound Library (MedChem Express, HY-L050), comprising 257 small molecule modulators targeting key enzymes in the ubiquitin pathway, was used for this screening. Data analysis was conducted in R, calculating the mean nuclei count for each treatment and comparing it to vehicle controls using t-tests (n=4). **(B)** The effect of UPS pathway inhibitors on bacterial growth in axenic culture was assessed at 24 hours post-infection (hpi), with compounds also used at a concentration of 10 µM. Data analysis in R involved calculating mean bacterial counts from triplicate samples, determining fold changes, and assessing statistical significance using t-tests, with results visualized in a volcano plot to highlight significant modulators.

The axenic growth assay, designed to evaluate the direct antimicrobial activity of compounds in a *Salmonella*-only culture (free of host cells), revealed that several tested compounds could inhibit bacterial growth. Notably, Bortezomib not only reduced the intracellular bacterial load but also directly inhibited *Salmonella* growth in axenic culture, reducing viability by at least 50%. Similarly, compounds such as Thiolutin and Ixazomib citrate demonstrated significant effects, reducing bacterial growth in both intracellular infection models and axenic cultures at a concentration of 10 μM (**Fig. 2B**). While these compounds hold promise as potential antimicrobials, their mechanisms of action might be independent of the UPS pathway, considering *Salmonella* lacks proteasomes. The antimicrobial properties of Thiolutin are well-established (26).

Those compounds that reduced intracellular bacteria without significantly affecting host cell viability (**Table 2**) included HBX 19818, OTUB1/USP8-IN-1, USP25/28 inhibitor AZ1, USP8-IN-3, VII-31, and others. These compounds were effective in enhancing bacterial clearance without affecting cell viability, and without affecting *Salmonella* viability in axenic culture conditions, making them promising candidates for further therapeutic development as host-directed antimicrobials.

**Table 2.**
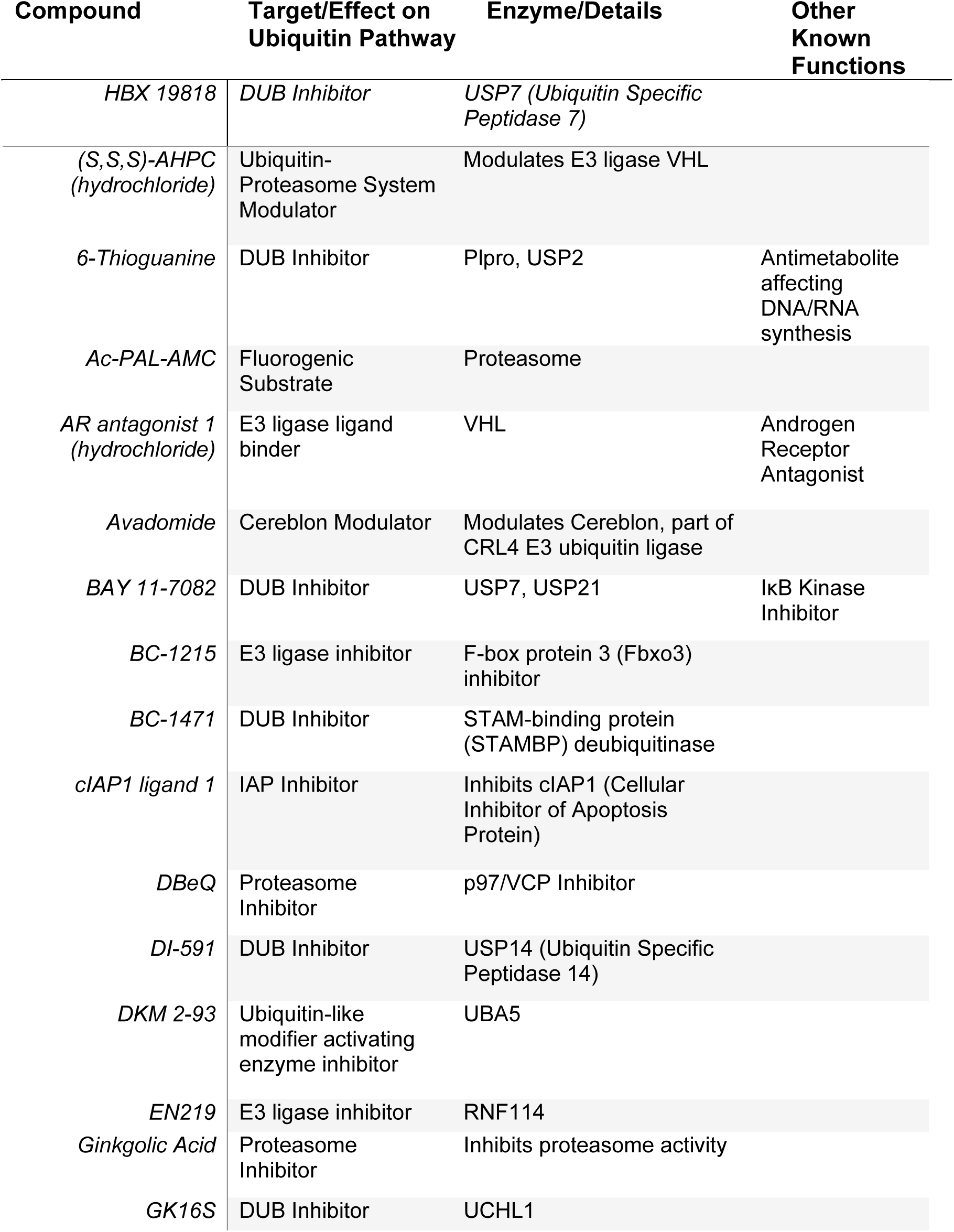
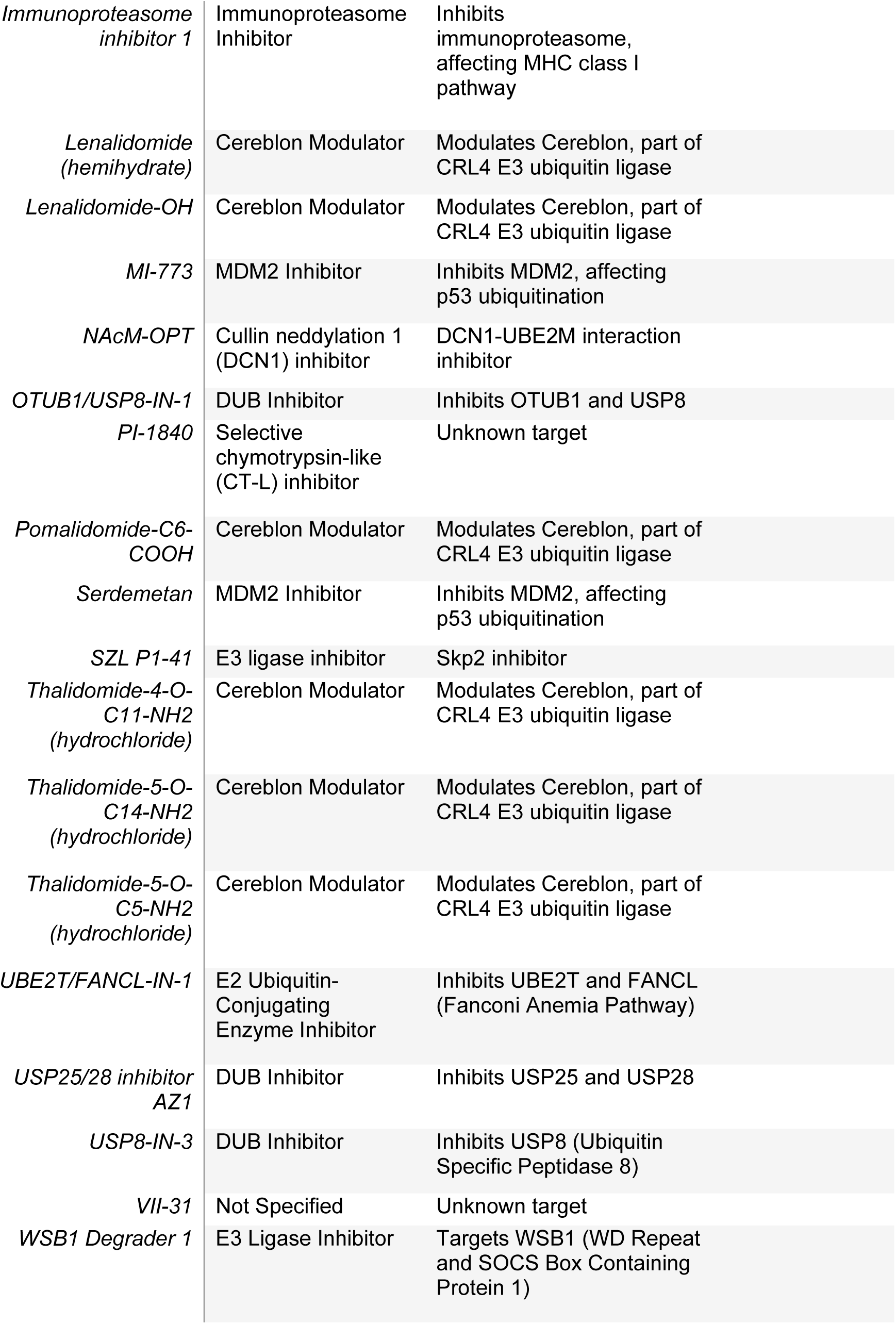
Ubiquitin pathway-modulating compounds with a significant effect on macrophage-mediated bacterial clearance without affecting viability, and their targets within the UPS and other functions.

### 2.2 Effect of the ubiquitin pathway inhibitors on TNF-α release from infected cells

Next, we evaluated the impact of selected DUB inhibitors and other UPS modulators on TNF-α secretion in RAW264.7 macrophages infected with *Salmonella* (**Fig. S1, Table S1**). TNF-α, a key pro-inflammatory cytokine released by infected macrophages, was quantified using ELISA at 24 hours post-infection (hpi). The vehicle control, infected with *Salmonella*, served as the baseline (100%) for comparing changes in TNF-α levels. A range of responses was observed, with some inhibitors significantly increasing TNF-α secretion, while others reduced it below baseline levels (**Fig. S1**). In total, 84 compounds significantly altered TNF-α secretion by at least 20%, either upregulating or downregulating its levels (**Table S1**), but 67 compounds were significant without altering cell viability (**Fig. S1**). Among the compounds that increased TNF-α, the USP7/USP47 inhibitor had the most pronounced effect, nearly tripling TNF-α levels. Similarly, USP18-IN-3, ML364, and HB007 almost doubled TNF-α levels, while USP9X-IN-1 resulted in a significant 30% increase. These compounds highlight potential pathways for enhancing inflammatory responses, which could be beneficial in specific therapeutic contexts but may also pose risks of physiological damage. Conversely, several compounds significantly decreased TNF-α levels. Notable examples include (S,R,S)-AHPC-Boc, which reduced TNF-α by 35%, ALV1 by 42%, and ALV2 by a substantial 53%. Additionally, AR antagonist 1 (hydrochloride) and Alloxan (hydrate) decreased TNF-α by 34% and 25%, respectively. These findings identify potential candidates for mitigating hyperinflammatory conditions. Interestingly, some compounds that modified TNF-α levels, such as AR antagonist 1 (hydrochloride), BC-1215, DBeQ, MI-773, Thalidomide-5-O-C14-NH2 (hydrochloride), and cIAP1 ligand 1, also demonstrated the capacity to reduce infection in RAW264.7 macrophages.

### 2.3 Alterations in DUB transcript levels post-*Salmonella*-infection

Since many of the UPS modulators were inhibitors of DUBs, we next analyzed the transcript levels of DUBs in RAW 264.7 macrophages following infection with the laboratory strain of *Salmonella* Typhimurium. Quantitative real-time PCR was employed to assess the expression changes in 70 DUBs between *Salmonella*-infected and non-infected RAW 264.7 macrophages at 2 and 24 hpi (**Fig. 3**). At the 2 hpi timepoint, three DUB transcripts were significantly upregulated, while seven were significantly downregulated (**Fig. 3A**). By 24 hpi, the number of significantly upregulated DUBs increased to ten, while 19 DUBs showed significant downregulation (**Fig. 3B**). At 2 hpi, *Usp2, Usp1*, and *Uchl5* were downregulated, while *Usp28*, *Usp24, Yod1, Otub1, Usp31, Usp36,* and *Usp18* were upregulated. Notably, DUBs such as *Otud1, Otud3, Otud7b, Uchl1, Usp11, Usp18, Usp22, Usp25, Usp27x, Usp31, Usp32, Usp36, Usp4, Usp46, Usp47, Usp49, Usp54*, and *Yod1* exhibited substantial increases in expression at 24 hpi, suggesting their potential involvement in the macrophage response to *Salmonella* infection. Conversely, DUBs including *Josd1, Otud4, Otud6b, Uchl5, Usp1, Usp14, Usp2, Usp36, Usp39*, and *Usp45* were among those downregulated at 24 hpi, indicating a potential suppression of their expression during infection. Interestingly, *Usp18, Usp31, Usp36*, and *Yod1* were consistently upregulated at both time points (2 and 24 hpi), suggesting a sustained role in the macrophage response to *Salmonella* infection. In contrast, *Usp1, Usp2*, and *Uchl5* remained downregulated across both time points, indicating a prolonged suppression during infection. To validate transcript-level findings, we examined the protein expression of one key DUB, USP25, at later infection times because protein synthesis often lags behind transcriptional changes. USP25 protein levels were significantly upregulated in infected macrophages by 26 hpi, with increases apparent at earlier time points (4 and 6 hpi) (**Fig. 3C, D**). These findings highlight the dynamic regulation of DUBs during *Salmonella* infection and suggest a critical role for specific DUBs, such as USP25, in the macrophage response.

**Figure 3.**
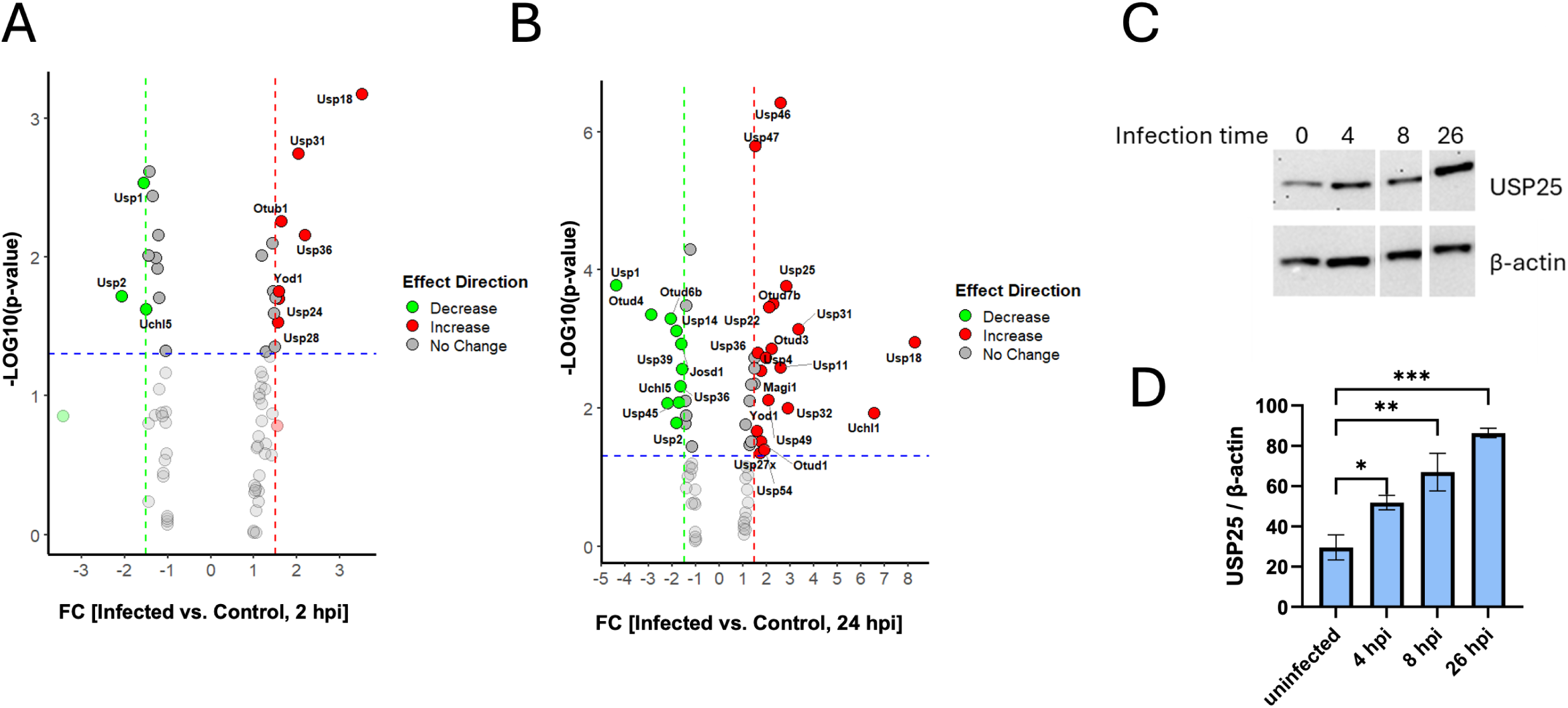
Regulation of DUB Expression During *Salmonella* Infection in Murine Macrophages. **(A-B)** Quantitative polymerase chain reaction (qPCR) analysis of deubiquitinating enzyme (DUB) transcript levels was performed using a Prime PCR array in RAW 264.7 macrophages infected with *Salmonella* Typhimurium strain UK-1. Panels (**A**) and (**B**) show the changes in DUB transcript levels at 2 hours and 24 hours post-infection, respectively, as analyzed in R. **(C)** USP25 expression in RAW 264.7 cells at 4, 8, and 26 hours post-infection (hpi) of RAW 264.7 cells with *Salmonella* (MOI 30:1) was performed using western blotting. (D). Protein levels were measured by ImageJ and normalized to beta-actin. The bars represent the relative protein levels of USP25.

Using published datasets, we analyzed the modulation of DUBs in monocyte-derived dendritic cells (moDCs) infected with three distinct *Salmonella* strains: Ty2 (*Salmonella* serovar Typhi), D23580 (*Salmonella* serovar Typhimurium, clinical isolate, invasive non-typhoidal *Salmonella* strain, iNTS), and LT2 (*Salmonella* serovar Typhimurium, laboratory strain, NTS) at 6 hpi (27). RNA sequencing of *Salmonella*-infected moDCs, sorted via fluorescence-activated cell sorting at 6 hpi, revealed significant differential gene expression between infected and uninfected cells in terms of DUB expression levels. At 6 hpi with the Ty2 strain, several DUB transcripts were significantly regulated. *Otud1, Otud7b, Usp11, Usp18*, and *Usp25* exhibited significant upregulation, while *Otud6b* and *Otud3* were downregulated, though *Otud3* was upregulated in our study (**Fig. S2A**). Similarly, infection with the D23580 strain resulted in the upregulation of *Otud1, Otud7b, Usp11, Usp18,* and *Usp25* at 6 hpi, while *Otud6b* was downregulated (**Fig. S2B**). In moDCs infected with the LT2 strain, *Otud1, Otud7b, Usp11, Usp18, Usp25*, and *Usp36* exhibited substantial increases in expression at 6 hpi (28). Conversely, DUBs such as *Otud6b, Usp39,* and *Usp45* were downregulated, along with Otud3, which was significantly upregulated in our study (**Fig. S2C**). A time-course RNA-seq analysis of DUB expression in LT2-infected moDCs at 2, 4, and 6 hpi (**Fig. S2D**) revealed that *Usp25, Usp18, Usp11*, and *Usp41* increased at later stages, suggesting roles in sustaining the immune response, while *Otud4* and *Uchl5* decreased, indicating early regulatory functions.

Collectively, the differential expression analysis of DUBs in response to diverse Salmonella strains—including typhoidal, non-typhoidal, and invasive strains (Ty2, D23580, and LT2)—as well as in RAW 264.7 macrophages infected with a laboratory strain of S. Typhimurium, revealed consistent regulatory patterns. Across all these studies, DUBs such as *Otud1, Otud7b, Usp11, Usp18*, and *Usp25* were significantly upregulated, while *Otud6b* was consistently downregulated.

### 2.4. Efficacy of AZ-1 in promoting clearance of intracellular *Salmonella*

The consistent upregulation of Usp25 in macrophages and moDCs during *Salmonella* infection, coupled with its identification in a high-throughput screen as a standout target, led us to evaluate AZ-1, a USP25/28 inhibitor. AZ-1 significantly reduced intracellular *Salmonella* without affecting host cell viability (**Fig. 1**). Its ability to enhance macrophage-mediated bacterial clearance consistently and safely highlights its strong potential for further development. In addition to AZ-1, we tested USP1 and UCH-L1 inhibitors, as these DUBs were differentially expressed during infection (**Fig. 4**).

**Figure 4.**
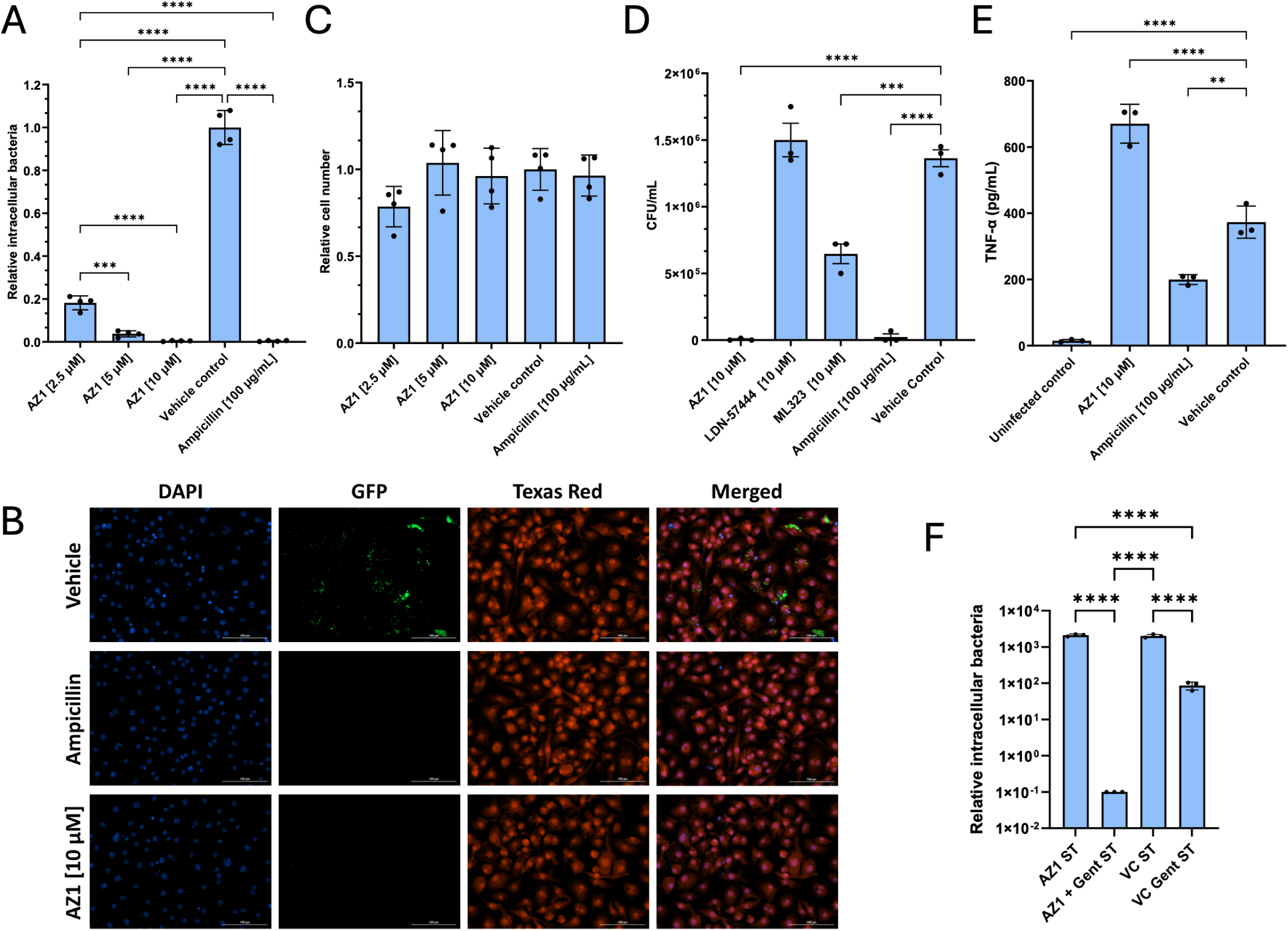
AZ-1 (USP25/28 inhibitor) effectively clears *Salmonella* in host cells. **(A-C) (A-C).** Representative images and quantification of *Salmonella* Typhimurium strain UK-1-infected macrophages (RAW 264.7) treated with AZ-1, ampicillin (positive control), or vehicle (DMSO). Ampicillin serves as a positive control for bacterial clearance. Infection was performed at a multiplicity of infection (MOI) of 30:1 for total of 26 hours. **(A).** Hoechst staining (blue) labels nuclei, while GFP fluorescence (green) indicates intracellular *Salmonella*. **(B).** Merged images show Hoechst (blue), GFP-Salmonella (green), and CellMask (red) to highlight cell structure. **(C).** Quantification of infected cells, comparing treatments. **(D).** Intracellular bacterial survival was quantified using a gentamicin protection assay followed by CFU counts, evaluating the effects of AZ-1, LDN-57444 (UCH-L1 inhibitor), ML-323 (USP1 inhibitor), and ampicillin (positive control), with the vehicle control serving as a negative reference. **(E).** RAW 264.7 cells were infected with *Salmonella* (MOI 30:1) and treated with AZ-1 or ampicillin (positive control), while the vehicle control served as a negative reference. TNF-α secretion levels were measured from culture supernatants 24 hours post-infection, and statistical analysis was conducted using one-way ANOVA followed by Tukey’s multiple comparisons test (n = 3). Error bars represent the mean ± SEM. **(F).** Gentamicin (100 μg/mL) was applied for 1 hour post-infection with *Salmonella* (MOI 30:1) of RAW 264.7 cells to eliminate extracellular bacteria, followed by its removal. Treatments for the remainder of the infection included AZ-1 alone (no gentamicin was added), AZ-1 combined with a lower concentration of gentamicin (30 μg/mL), vehicle control, and vehicle control (DMSo) with a lower concentration of gentamicin (30 μg/mL). Hoechst staining (blue) labeled nuclei, GFP fluorescence indicated intracellular *Salmonella*, and CellMask highlighted cytoplasm to enable intracellular bacterial count. Quantification compared bacterial survival across conditions, evaluating the effects of AZ-1 with or without gentamicin treatment.

AZ-1 significantly reduced intracellular bacterial counts in a dose-dependent manner, with maximal activity observed at 10 μM, comparable to the efficacy of ampicillin (**Fig. 4A, B**). Importantly, cell viability remained unaffected at all concentrations tested (**Fig. 4C**), and no cytotoxicity detected in uninfected cells treated with AZ-1 (**Fig. S3**). Colony plating further confirmed that AZ-1 significantly reduced intracellular bacterial counts compared to vehicle controls and other tested DUB inhibitors (**Fig. 4D**). In addition to its antimicrobial effects, AZ-1 also significantly upregulated TNF-α secretion during *Salmonella* infection (**Fig. 4E**), suggesting it may modulate the immune response in a beneficial way.

To investigate the role of gentamicin in facilitating AZ-1’s intracellular activity, we conducted an experiment where RAW 264.7 macrophages were infected with *Salmonella* Typhimurium for 1 hour. After the infection, extracellular bacteria were eliminated by applying gentamicin (100 μg/mL) for an additional hour, followed by its removal. Cells were then treated for the remainder of the infection period with either AZ-1 alone, AZ-1 combined with a lower concentration of gentamicin (30 μg/mL), vehicle control, or vehicle control with gentamicin. The results showed that AZ-1 alone did not reduce the intramacrophage bacterial load in the absence of gentamicin, while the combination of AZ-1 with gentamicin led to a significant 4-log reduction in bacterial counts (**Fig. 5F**). This suggests that AZ-1’s ability to clear intracellular *Salmonella* is dependent on the effective suppression of extracellular bacteria by gentamicin. To exclude the possibility that gentamicin alone contributes to intracellular bacterial clearance over prolonged periods, we evaluated various sequential treatment regimens after 1 hour infection with *Salmonella* of RAW 264.7 macrophages. Specifically, we used an initial high concentration of gentamicin (applied for 1 hour) to eliminate extracellular bacteria, followed by lower sustained concentrations during the overnight incubation. The tested regimens included the following initial/sustained gentamicin concentration pairs (in μg/mL): 100/25, 50/5, 25/2.5, and 100/20. These regimens were combined with either vehicle control or AZ-1 treatment. In all conditions, AZ-1 consistently reduced bacterial load after overnight infection, reinforcing its efficacy when extracellular bacteria are effectively controlled by gentamicin (**Fig. S4**). By testing multiple gentamicin dosing strategies, we confirmed that prolonged gentamicin exposure alone does not significantly influence the clearance of intracellular bacteria. Instead, the observed reduction in bacterial loads is attributed to AZ-1’s intracellular activity, which depends on the suppression of extracellular bacteria during the initial infection period.

**Figure 5.**
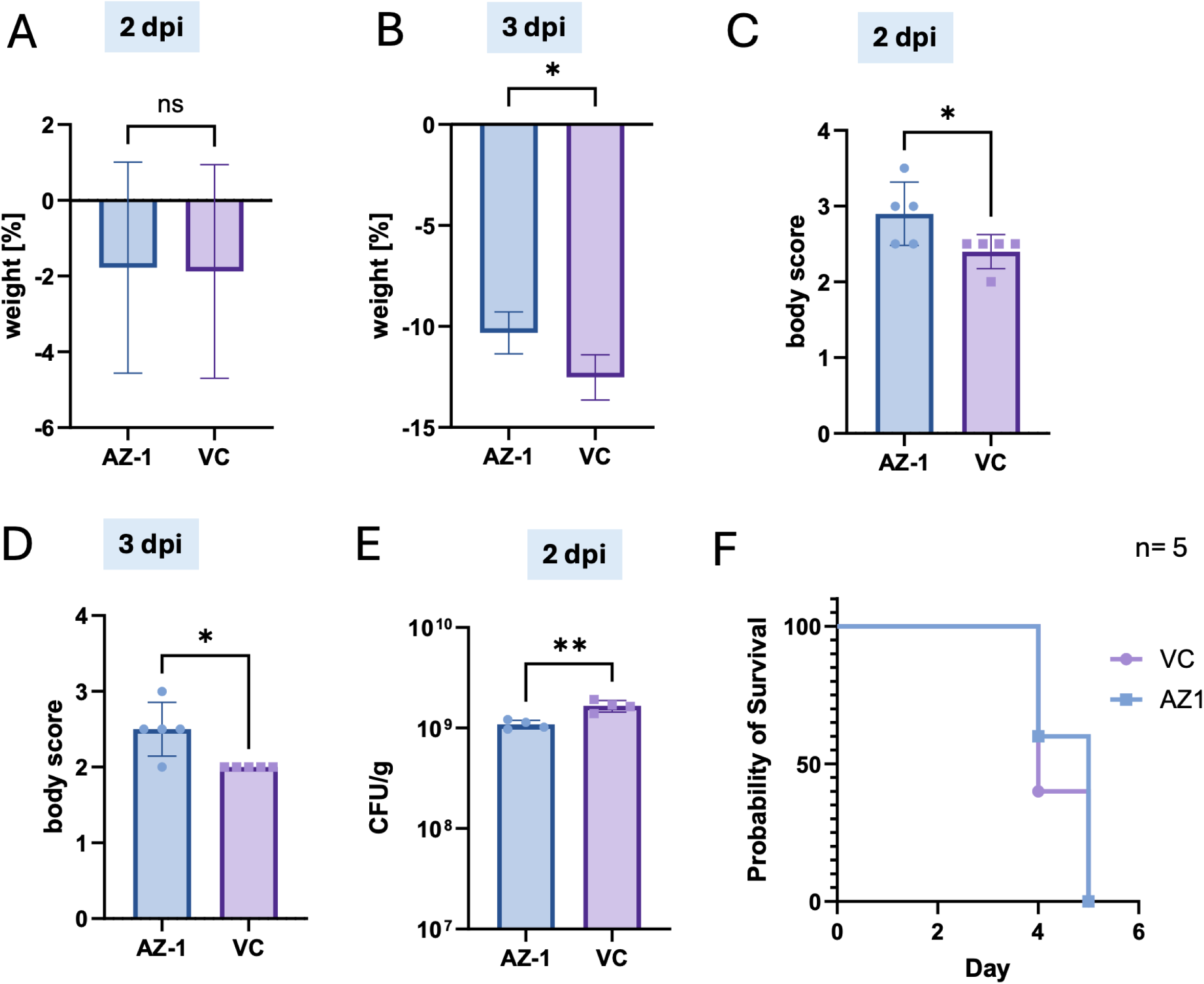
Treatment of C57BL/6J mice with AZ-1 influences weight loss and moderately reduces fecal bacterial load, but does not prolong survival administered alone. **(A-C).** Mice infected with *Salmonella* were treated with AZ-1 (20 mg/kg) or vehicle control (VC) starting on day 1 post-infection. Body weight was measured daily to assess infection severity. Weight loss recorded on day 2 **(A**), day 3 **(B)**, post-infection are shown. **(C-D).** Body scores were collected at 2 and dpi. **(E).** Fecal bacterial loads were quantified 2 dpi by plating stool samples on selective agar and counting colony-forming units (CFUs). **(F).** Survival of *Salmonella*-infected mice treated with AZ-1 or vehicle control (DMSO) was monitored over the course of the experiment using Kaplan-Meier analysis. Data represent mean ± SEM. Statistical significance was determined using a two-tailed t-test for body weight loss and CFU data, and a log-rank test for survival analysis. *p < 0.05.

These findings highlight the potential of AZ-1 as part of a combination therapy for *Salmonella* infections. By promoting macrophage-mediated bacterial clearance, preserving cell viability, and enhancing TNF-α production, AZ-1 may serve as a valuable adjunct to conventional treatments like gentamicin for managing bacterial infections. AZ-1 specifically appears to limit intramacrophage growth of *Salmonella* through currently unknown mechanisms.

### 2.5. *In vivo* evaluation of AZ-1 reveals partial efficacy in reducing *Salmonella* infection severity in a murine model

We next evaluated the *in vivo* antimicrobial efficacy of AZ-1, identified during the primary screening phase, using a streptomycin-pretreated murine model of *Salmonella* infection. Streptomycin was administered to disrupt the gut microbiota and facilitate *Salmonella* colonization the following day (29), providing a good model for assessing bacterial burden and treatment effects. Mice were treated with AZ-1 (20 mg/kg) or vehicle control (DMSO) starting on day 1 post-infection. At 2 days post-infection (dpi), there was no significant difference in weight loss between AZ-1-treated and vehicle-treated mice (**Fig. 5A**). However, by 3 dpi, mice treated with AZ-1 exhibited significantly reduced weight loss compared to vehicle controls (**Fig. 5B**). Similarly, body condition scores were significantly higher for AZ-1-treated animals at both 2 and 3 dpi compared to controls (**Figs. 5C, D**). Additionally, AZ-1 treatment significantly reduced fecal bacterial loads on day 2, as determined by CFUs (**Fig. 5E**), suggesting initial enhanced bacterial clearance from the gastrointestinal tract or reduced colonization during early infection.Despite these improvements in clinical parameters, survival analysis revealed no significant difference in survival rates between AZ-1-treated and vehicle-treated mice (**Fig. 5F**). While AZ-1 alleviates aspects of disease severity, it does not independently improve survival under the conditions tested.

These findings indicate that AZ-1 has the potential to mitigate infection severity and reduce bacterial burden but may require combination with other therapies to maximize its therapeutic effectiveness against *Salmonella* infections, consistent with our in vitro data (**Fig. 4F**).

### 2.6. Effect of AZ-1 inhibitor on the MAPK and NF-κB signaling

*Salmonella* infection is associated with alterations in host signaling pathways, including ERK1/2 (30) and NF-κB, which are key regulators of immune responses and inflammation. Several *Salmonella* virulence factors are known to interfere with the NF-κB pathway (31) (32) (33). Understanding how these pathways are modulated during infection, as well as how potential therapeutic agents like AZ-1 influence these dynamics, could uncover novel mechanisms to mitigate bacterial pathogenesis. We analyzed the temporal dynamics of TRAF3 expression, ERK1/2 activation, and NF-κB signaling during *Salmonella* infection with or without AZ-1 treatment. TRAF3, a known negative regulator of the non-canonical NF-κB pathway, was upregulated during the early stages of infection (4 hpi) in response to *Salmonella* infection (**Fig. 6A, B**). In contrast, AZ-1 treatment significantly enhanced TRAF3 levels during the late stages of infection (26 hpi, 24 hours post-AZ-1 treatment). Additionally, AZ-1 treatment reduced the p-ERK1/2 to ERK1/2 ratio at 26 hpi compared to vehicle-treated infected cells, indicating that AZ-1 attenuates ERK1/2 activation during the late stages of *Salmonella* infection. However, no significant effects were observed at earlier time points (**Fig. 6A, C**).

**Figure 6.**
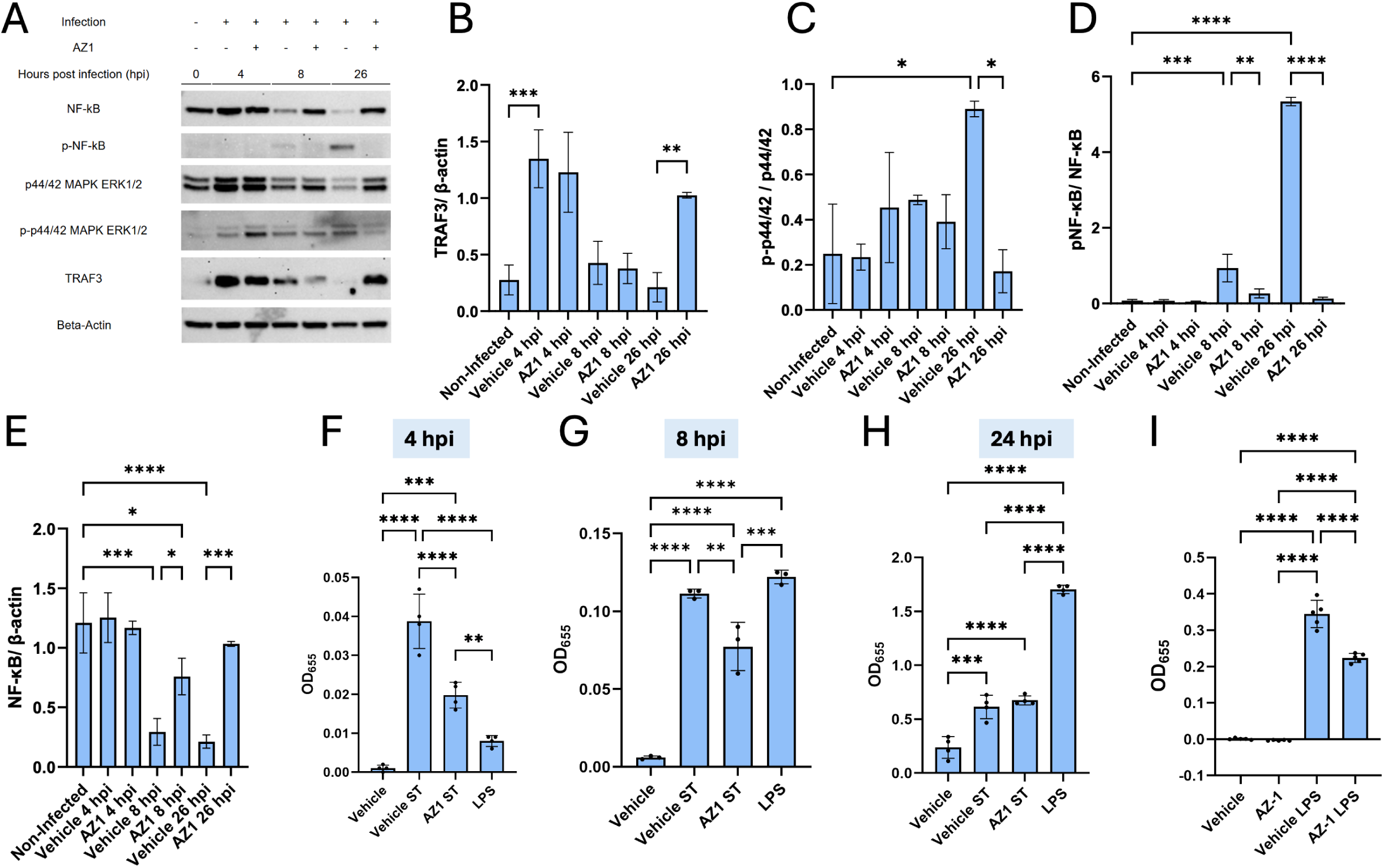
AZ-1 affect NF-kB signaling. **(A-E).** RAW264.7 macrophages were infected with UK-1 *S.* Typhimurium at MOI 30:1 and incubated for 1 hour. Following this period, cells were treated with 100 µg/mL gentamicin for an additional hour under the same conditions to eliminate extracellular bacteria. After the gentamicin incubation, the media was aspirated, cells were washed, and fresh media containing either AZ-1 or vehicle control (DMSO) was added. Cell lysates were collected at 4, 8, and 26 hours post-infection (hpi), along with an uninfected 0-hour baseline control, and prepared for Western blot analysis **(A).** A Western blot was performed using antibodies specific for NF-κB, phosphorylated NF-κB (p-NF-κB), p44/42 MAPK ERK1/2, p-p44/42 MAPK ERK1/2, TRAF3, USP25, and β-actin. ImageJ was used to measure the relative band intensity for each of these antibodies across three biological replicates for each condition with ratios performed for TRAF3 / β-actin **(B)**, p-p44/42 / p44/42 **(C)**, p-NF-κB / NF-κB **(D)**, and NF-κB / β-actin **(E)**. (**F-H**). Aa parallel infection model was performed using InvivoGen’s RAW 264.7-Blue NF-κB reporter cell line. Supernatants were collected at 4 **(F)**, 8 **(G)**, and 26 hpi **(H)** to measure NF-κB-driven secreted alkaline phosphatase (SEAP) activity. **(I).** RAW 264.7-Blue cells were treated with 500 ng/mL lipopolysaccharide (LPS) for 6 hours in the presence of either a vehicle control in DMSO or AZ1 (I).

*Salmonella* infection activated NF-κB signaling, as demonstrated by increased phosphorylation of NF-κB (p-NF-κB) in western blots at both 8 hpi and 26 hpi (**Fig. 6A, D**). Importantly, AZ-1 treatment markedly downregulated NF-κB phosphorylation at these time points, suggesting diminished NF-κB activation. Interestingly, the total NF-κB expression was increased in AZ-1-treated cells at 8 hpi and 24 hpi (**Fig. 6A, F**). The effect of AZ-1 on NF-κB activation was quantitatively validated using the Quanti-Blue assay, which measures NF-κB-driven secretion of alkaline phosphatase (SEAP) through colorimetric OD655 readings. AZ-1-treated cells exhibited significantly reduced NF-κB activity compared to vehicle-treated controls at 4 hpi and 8 hpi, aligning with the western blot findings. However, these differences were not observed at 24 hpi using this readout (**Fig. 6F, G, H**). A similar trend was observed in LPS-treated cells, where AZ-1 also decreased LPS-induced NF-κB activation following 6 hours of LPS treatment (**Fig. 6I**).

In conclusion, AZ-1 treatment modulates NF-κB and ERK1/2 signaling during *Salmonella* infection, highlighting a role for AZ-1 in targeting host immune signaling pathways, potentially contributing to enhanced bacterial clearance.

### 2.7 Efficacy of AZ-1 in promoting clearance of other bacterial infections

Subsequently, we tested the efficacy of AZ-1 on the clearance of other pathogens, including ESKAPE pathogens. ESKAPE pathogens—*Enterococcus faecium, Staphylococcus aureus, Klebsiella pneumoniae, Acinetobacter baumannii, Pseudomonas aeruginosa*, and *Enterobacter* species—are a group of antibiotic-resistant bacteria known for causing severe hospital-acquired infections (34). RAW 264.7 cells were infected with *P. aeruginosa* PAO1*, K. pneumoniae* KPPR1, and *A. baumannii* MAB103 at MOI of 10. The infected macrophages were treated with AZ-1 or DMSO (control). After 4 hours of treatment, the cells were lysed, and the bacterial load was determined by plating the lysates for CFU enumeration. The results demonstrate a significant reduction in CFUs for all three tested ESKAPE pathogens in the AZ-1-treated samples compared to the control, indicating the efficacy of AZ-1 in promoting clearance of the ESKAPE pathogens (**Fig. 7**). Testing AZ-1 and HBX 19818 on the intramacrophage growth of another Gram-negative bacterium, *Francisella novicida*, which has a different intracellular llifestyle, evaluated using immortalized murine bone marrow-derived macrophages and RAW 264.7 cells indicatde that AZ1 and HBX 19818 had no effect on the growth of *F. novicida* within macrophages in the tested condutions (**Fig. S5**).

**Figure 7.**
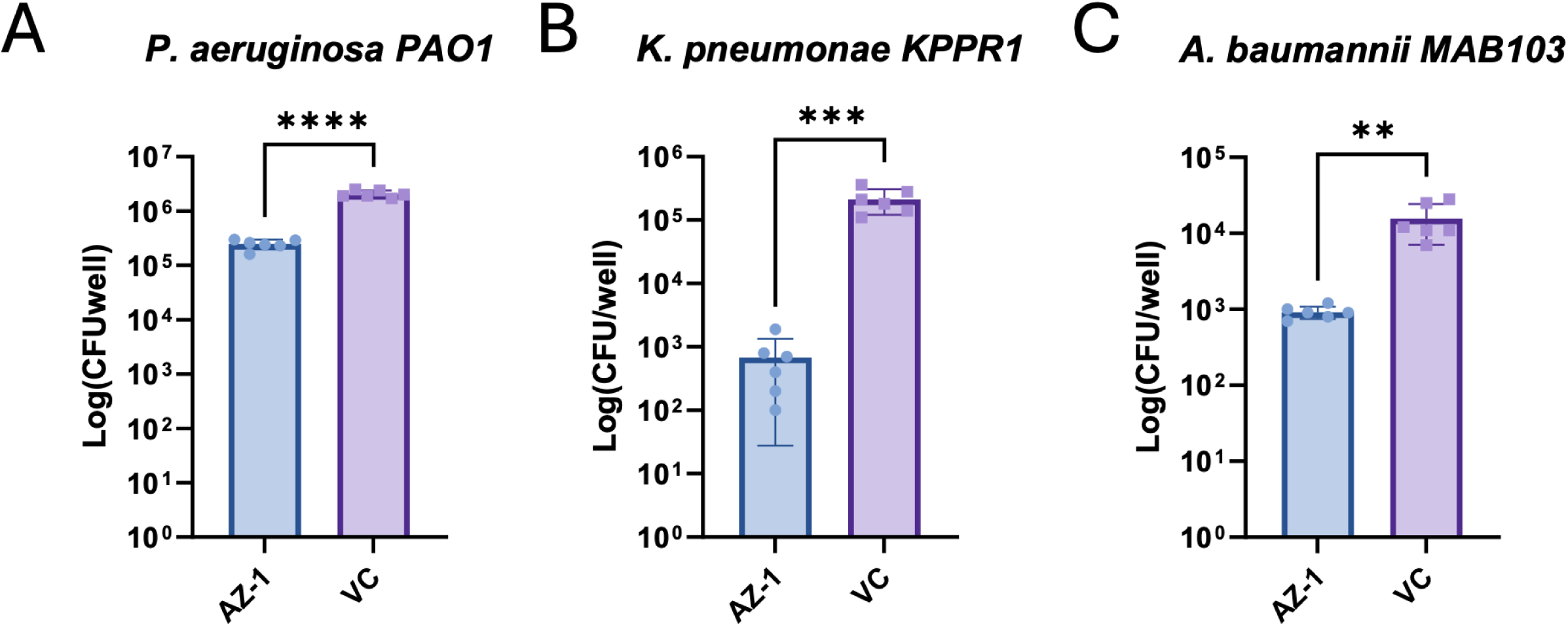
Effects of AZ1 on the intramacrophage growth of various ESKAPE pathogens. RAW 264.7 cells were infected with *Pseudomonas aeruginosa, Klebsiella pneumonia*e, and *Acinetobacter baumannii* at a multiplicity of infection (MOI) of 10. After washing and gentamicin treatment to remove extracellular bacteria, the infected macrophages were treated with 10 μM AZ1 or DMSO vehicle (VC) control. The cells were lysed, and the bacterial load was assessed by plating the lysates for colony-forming unit (CFU) enumeration. Bars represent the mean CFUs ± standard error of the mean (SEM). N=6. The experiment was repeated three times, and a representative graph is shown. Statistical significance between treatments was determined using Student’s t-test (****p < 0.0001)

These findings show the importance of testing such novel host-targeting inhibitors across various bacterial pathogens and demonstrate the potential of the tested drugs as therapeutic agents for infections caused by *Salmonella*, *Pseudomonas*, *Klebsiella*, and *Acinetobacter*, but not *Francisella*.

## 3. Discussion

The escalating crisis of antibiotic resistance among bacterial pathogens necessitates innovative therapeutic approaches. This study demonstrates the potential of host-targeted therapeutics (HTTs) to combat infections by focusing on DUBs and their role in macrophage responses to *Salmonella*. Targeting host pathways rather than pathogens directly offers distinct advantages, particularly against bacteria like *Salmonella* and ESKAPE pathogens, which frequently resist conventional antibiotics (34).

High-throughput screening of a UPS modulator compound library revealed several promising candidates that significantly enhance bacterial clearance and modulate inflammatory responses without compromising host cell viability. Among them, DUB inhibitors were particularly effective, targeting key enzymes like USP7, USP8, and USP25 to disrupt Salmonella’s immune evasion mechanisms. Proteasome inhibitors such as DBeQ and ginkgolic acid (GA) also exhibited significant activity, which are known to be acting through ubiquitin-dependent and autophagic pathways (35–40). Additionally, CRBN modulators, including avadomide and lenalidomide derivatives, here reduced bacterial loads likely by targeting the CRL4 E3 ubiquitin ligase complex (41). MDM2 inhibitors (e.g., MI-773, serdemetan) effectively combat intracellular *Salmonella* by modulating the MDM2-p53 axis, critical for host defense(42–47). Beyond *Salmonella*, MDM2 is important in controlling outcomes of other infections, such as *Chlamydia* (48)and *Helicobacter pylori*, where it regulates p53-mediated host responses (49). The potential to disrupt Salmonella effector AvrA-mediated p53 stabilization presents a novel therapeutic avenue (50). Evaluating MDM2 inhibitors’ impact on AvrA-mediated virulence and p53 stabilization could provide new therapeutic strategies. Similarly, neddylation inhibitors (e.g., NAcM-OPT, VII-31) modulate immune responses by influencing cytokine production and inflammatory signaling in macrophages and neutrophils. Their ability to reduce pro-inflammatory cytokines, such as TNF-α and IL-6, and regulate the NF-κB pathway highlights their dual potential in limiting inflammation and enhancing host defenses (51) (52, 53) (54).

Several DUB inhibitors showed marked efficacy in clearing intracellular *Salmonella* infections, including These include HBX 19818, 6-Thioguanine, BAY 11-7082, BC-1471, DI-591, GK16S, OTUB1/USP8-IN-1, the USP25/USP28 inhibitor AZ1, and USP8-IN-3. These inhibitors target key enzymes within the host’s ubiquitin-proteasome system, including USP7, USP2, USP21, STAMBP, USP14, UCHL1, OTUB1, USP8, and USP25/USP28. For instance, USP8 inhibition has been linked to autophagy modulation and enhanced *Salmonella* clearance(17). USP7 plays a role in inflammasome regulation and NF-κB stabilization, with its inhibition significantly increasing TNF-alpha levels in our dataset, highlighting its complex role in infection (55). USP7 inhibition by three different inhibitors resulted in a significant increase in TNF-alpha levels following *Salmonella* infection. Other DUBs like UCH-L1 and OTUB1 influence bacterial invasion by modulating actin cytoskeleton dynamics and autophagy. For instance, UCH-L1 inhibition reduces *Listeria* and *Salmonella* uptake(56), while OTUB1 knockdown decreases *Yersinia* infections through its effects on RhoA dynamics (57). However, limitations of current inhibitors, such as LDN-57444’s transient activity, underscore the need for more robust alternatives (58).

USP25, which is consistently upregulated during *Salmonella* infection, emerged as a promising target for host-directed therapies in our screen. AZ-1, a USP25/USP28 inhibitor, showed broad-spectrum potential by reducing bacterial loads of *P. aeruginosa, K. pneumoniae*, and *A. baumannii*. While these pathogens are typically classified as extracellular, they are known to have intracellular phases, particularly within macrophages, demonstrating their ability to exploit intracellular niches during infection (59–61). While AZ-1 was effective against *Salmonella* and the aforementioned pathogens, it showed limited efficacy against *F. novicida*. This may be due to *F. novicida*’s resistance to oxidative burst mechanisms (62, 63), although it remains speculative. Interestingly, recent studies have shown that USP25 is also induced during *Mycobacterium tuberculosis* infection, where it promotes antimycobacterial responses in macrophages, as demonstrated using USP25 knockout models (64). However, the application of AZ-1 in this context has not yet been explored. Future studies will aim to refine experimental conditions and further assess AZ-1’s therapeutic potential across different infection models.

It is currently not known how AZ-1 exerts it therapeutic function in the tested models. AZ-1 treatment does modulate NF-κB signaling as AZ-1 treamtent is associated with stabilizing unphosphorylated NF-κB at later stages of infection, while suppressing its activation early on, highlighting its potential to fine-tune inflammatory responses. *Salmonella* infection activates key pathways like ERK1/2 (30) and NF-κB, which influence bacterial survival in macrophages. ERK1/2 activation promotes COX-2 expression, supporting *Salmonella* survival via the PKA pathway (30), while NF-κB upregulates pro-survival and inflammatory responses. Previous studies have shown that USP25 deficiency increases p-ERK1/2 and MAPK activation(65), although minimal effects on MAPK signaling have been noted in IL-17-stimulated macrophages (66). In our study, AZ-1 stabilized TRAF3 during the later stages of *Salmonella* infection, suggesting a delayed regulatory role for USP25. Interestingly, AZ-1 suppressed NF-κB activation at early time points by reducing phosphorylation and activity, but this effect diminished at later stages (26 hpi). Notably, AZ-1 impacted the stability of unphosphorylated NF-κB at both 8 and 26 hpi, demonstrating its potential to modulate temporal dynamics of this pathway. Although the precise mechanism of AZ-1’s enhancement of bacterial clearance remains unclear, its significant influence on the NF-κB pathway points to its role in this process.

Despite the promise of DUB inhibitors like AZ-1, the lack of specific inhibitors for many DUBs remains a challenge. Upregulated DUBs during *Salmonella* infection, such as USP11 (67), OTUD7B (68, 69) (70), and OTUD1, which are involved in innate immunity, may present future therapeutic targets. Conversely, downregulated DUBs like OTUD4 (71) and UCH-L5 (72) may indicate disruptions in host defense mechanisms that warrant further exploration.

In conclusion, while macrophages play a central role in immune defense, they can also act as ‘Trojan horses,’ aiding the systemic spread of intracellular pathogens like *Salmonella* (6–9). Our study highlights that targeting DUBs, particularly USP25, with inhibitors such as AZ-1 not only reduces bacterial load, likely disrupting the conditions that support *Salmonella* persistence within macrophages. However, our *in vivo* experiment indicated only a moderate effect of AZ-1 on bacterial colonization and weight loss during infection. These findings suggest that, in this proof-of-principle study, AZ-1 alone was insufficient to clear the infection. This is further supported by in vitro data showing that antibiotics targeting extracellular bacteria may need to be used in conjunction with such host-targeted therapies for optimal efficacy.

Beyond validating AZ-1, we identified a diverse catalog of UPS modulators that may serve as promising candidates for combating intracellular bacterial infections. The broader efficacy of AZ-1 against *Salmonella* and select ESKAPE pathogens highlights the potential of HTTs to complement traditional antibiotics. Combination strategies, pairing DUB inhibitors like AZ-1 with antibiotics, may address the limitations of single-drug therapies by targeting both bacterial survival within macrophages and extracellular pathogens. Antibiotics often struggle to penetrate intracellular compartments (73), making combination therapies a compelling strategy to achieve effective clearance of persistent intracellular pathogens. Future research should focus on elucidating the mechanisms of DUB regulation during infection, optimizing combination therapies, and evaluating their preclinical efficacy. By addressing the growing threat of antibiotic resistance, these strategies offer a promising avenue for advancing treatment options and mitigating the global burden of intracellular infections.

## 4. Methods

### 4.1 Cell and *Salmonella* cultures

RAW264.7 macrophages, a murine macrophage cell line, were cultured in Dulbecco’s Modified Eagle Medium (DMEM) supplemented with 10% fetal bovine serum (FBS) and 1% penicillin-streptomycin.

The overnight cultures of *Salmonella enterica* serovar Typhimurium, strain 12023 or UK-1 c3761, were cultivated at 37°C in lysogeny broth (LB) Miller media overnight with shaking as we previously described (74, 75). Overnight bacterial cultures were diluted in fresh LB Miller media in a 1:80 ratio to reach an optical density at 600 nm (OD_600_) of 0.05 and grown until OD_600_ was 0.50 for three hours, followed by washing with PBS and resuspension in cell culture medium. GFP-expressing *Salmonella* UK-1 was generated by electroporation with pON::sfGFP plasmid as previously described (76).

### 4.2 High-throughput screening of ubiquitination pathway modulators for enhancing macrophage-mediate bacterial clearance

The high-throughput screening (HTS) of ubiquitination pathway modulators involved a systematic process designed to evaluate the efficacy of various compounds in enhancing bacterial clearance in *Salmonella*-infected macrophages (**Fig. 1)**. The largest groups of compounds in our library targeted the ubiquitination pathway, with 75 aimed at E1/E2/E3 enzymes and 59 targeting DUBs. Additionally, 58 compounds were ligands for E3 ligases, 52 targeted apoptosis, and 50 targeted the proteasome. The MDM-2/p53 interaction, key in immune response and cellular stress, was targeted by 26 compounds, and autophagy by 16 compounds. Other significant categories included molecular glues (14 compounds), p97 inhibitors (11 compounds), and NF-κB pathway inhibitors (9 compounds). We also included compounds targeting specific proteins and pathways involved in macrophage function, such as Bcl-2, Akt, IKK, and cathepsin. Smaller groups of compounds targeted DNA methyltransferases, caspases, oxidative phosphorylation, reactive oxygen species, PARP, and others, crucial for macrophage activation and bacterial clearance.

Initially, GFP-labeled *Salmonella* was cultured overnight in lysogeny broth (LB) Miller at 37°C with shaking. The bacterial culture was diluted in fresh LB to an OD600 of 0.05 and grown until the mid-log phase (OD_600_ =∼0.50). Concurrently, RAW264.7 macrophages were seeded into 96-well plates and allowed to adhere overnight. These macrophages were then infected with *Salmonella* at an MOI of 30:1. Following infection, the macrophages were treated with a library of DUB inhibitors at various concentrations to evaluate their effects on bacterial clearance. After treatment, the infected macrophages were incubated for a specified duration (2 or 24 hours), followed by two washes with Dulbecco’s PBS with calcium and magnesium (DPBS) with a subsequent fixing step using 4% paraformaldehyde (PFA). After the fixing step, the cells were washed twice with DPBS and permeabilized using a 0.1% Triton® X-100 solution in PBS. After the permeabilization step, the cells were washed twice with DPBS and stained with Hoechst dye and HCS Cell Mask Red to facilitate the assessment of cell viability number and bacterial load. The infected and treated macrophages were then imaged using a high-content imaging system (Cytation 5, Agilent), capturing GFP fluorescence from *Salmonella* and, Hoechst-stained nuclei, and Texas Red fluorescent cytoplasm. This imaging allowed for the quantification of intracellular bacterial load, normalized to cell viability number. The efficacy of each DUB inhibitor was determined by comparing bacterial loads in treated versus untreated control wells, with results visualized through fluorescence imaging. This HTS approach enabled the identification of promising inhibitors which significantly reduced intracellular bacterial loads in infected macrophages. To evaluate the efficacy of DUB inhibitors on bacterial clearance, we performed a statistical analysis using *R* studio (ver. 4.3.0). Libraries including *ggplot2*, *dplyr*, and *ggrepel* for calculations and data visualization. The analysis involved calculating mean values of replicates, performing t-tests to determine p-values, and computing fold changes relative to vehicle control. Significant results were visualized using a volcano plot, which displayed the log10 fold change against the -log10 p-value.

To evaluate the cell number after infection, we used DAPI counts of nuclei as our data source. The analysis involved processing the data to calculate average nuclei counts for each condition. The percentage of the cell number was then calculated relative to the Vehicle Control. Statistical significance was assessed using t-tests, and the results were visualized with a volcano plot created using R libraries *tidyverse*, *ggplot2*, and *ggrepel*. This plot displayed the percentage viability on the x-axis and the -log10(p-value) on the y-axis, highlighting significant changes in cell number.

### 4.3 *Salmonella* infection studies

RAW 264.7 macrophages were infected with GFP-labeled or wild-type *Salmonella* at MOI of 30:1. After 1 hour, gentamicin was added at a concentration of 100 ug/mL for 1 hour, washed with 1X DPBS, and lowered to 25 ug/mL gentamicin for the remaining amount of time. For the infections that included inhibitor studies, cells were infected for 1 hour, washed with PBS, and then media was added that included inhibitors at indicated concentrations (in the absence of gentamicin), including AZ-1, LDN-57444, and ML323, to evaluate their impact on bacterial clearance. Appropriate vehicle controls were used at the same volumetric concentration of DMSO as the inhibitor-containing stocks.

### 4.4 Intramacrophage *Francisella novicida* growth assay

Infections of murine bone marrow-derived macrophages (BEI resources, NR-9456) were performed as previously described (77). Briefly, 4.5 × 10^4 cells were seeded in the absence of antibiotics, washed, and exposed to *Francisella novicida* at MOI of 0.1. Plates were centrifuged at 1,000 × g for 30 minutes at room temperature, followed by incubation at 37°C for 30 minutes to ensure infection. Cells were then treated with 50 μg/mL gentamicin for 30 minutes to remove extracellular bacteria and washed thoroughly. Infected cells were subsequently treated with the indicated concentrations of the drugs, and cells were lysed immediately and 18 hpi by the addition of Triton X-100 to a final concentration of 0.1%. Lysates were diluted in tryptic soy broth + 0.1% cysteine and plated for colony-forming unit enumeration. Experiments were performed with at least four technical and three biological replicates.

### 4.5 Evaluation of AZ1 on intramacrophage growth of ESKAPE pathogens

RAW 264.7 cells were maintained in Dulbecco’s Modified Eagle’s Medium (GenClone 25-500) supplemented with 10% FBS and 1% Pen Strep (GenClone, 25-512) in a 37°C humidified incubator with 5% CO2. Two days prior to infection, cells were scraped, washed, and seeded in 96-well plates at a density of 5×10^4 cells/well in 100 μL of DMEM supplemented with 10% FBS without antibiotics. The day before infection, *P. aeruginosa* PAO1*, K. pneumoniae* KPPR1, and *A. baumannii* MAB103 were grown in Lennox Broth (LB) in a 37°C incubator shaking at 220 revolutions per minute (RPM). On the day of infection, eukaryotic cells in a duplicate plate were detached with trypsin at 37°C for 10 minutes and counted. Bacterial cultures were adjusted to OD600 = 1, washed three times by centrifugation at 10,000 x g for 10 minutes, and resuspended in phosphate-buffered saline (PBS). To achieve an MOI of 10, appropriate volumes of each bacterial suspension were diluted in PBS and added to DMEM supplemented with 10% FBS. Eukaryotic cells were washed with PBS, and 100 µL of the bacterial suspension was added to each well. Because *P. aeruginosa, K. pneumoniae,* and *A. baumannii* are extracellular bacteria, the infection was facilitated by centrifugation at 500 x g for 30 minutes at 37°C. The plate was then incubated for an additional 30 minutes in a humidified incubator with 5% CO2 at 37°C to allow the infection to progress for a total of one hour. After incubation, the media were removed, and wells were washed twice with PBS. One hundred microliters of DMEM supplemented with 10% FBS and 100 μg/mL gentamicin was added to each well to eliminate extracellular bacteria, and cells were incubated for one hour in a humidified incubator with 5% CO2 at 37°C. Following this incubation, the media was removed, wells were washed twice with PBS, and 100 μL of DMEM supplemented with 10% FBS, 25 μg/mL gentamicin, and either A) 10 μM AZ1 (1:1000 dilution of 10 mM dissolved in DMSO) or B) 1:1000 dilution of DMSO (control) was added to each well. Cells were then incubated for four hours in a humidified incubator with 5% CO2 at 37°C. After incubation, wells were washed twice with PBS, 100 μL of 0.05% Triton X-100 was added to each well and incubated in a humidified incubator with 5% CO2 at 37°C for 12 minutes. The resulting lysate was serially diluted, and 10 μL from each well was plated onto LB agar plates. Plates were incubated overnight at 37°C, and colony-forming units (CFUs) were enumerated.

### 4.6 qPCR analysis

The quantitative polymerase chain reaction (qPCR) analysis for assessing the transcript levels of murine DUBs was done for measurement of gene expression of DUBs during Salmonella infection. We designed a primer array of all the characterized murine DUBs to evaluate the expression of these enzymes during infection. *Salmonella* was cultured overnight in LB at 37°C with shaking. Parallel to this, RAW264.7 macrophages were seeded into culture dishes and allowed to adhere overnight. These cells were subsequently infected with the prepared *Salmonella* culture at an MOI of 30:1 as described in the sections above. Following the incubation period, the infected macrophages were lysed to extract total RNA. The isolated RNA was then subjected to reverse transcription to synthesize complementary DNA (cDNA), utilizing the iScript Reverse Transcriptase Supermix for RT-qPCR per the manufacturer’s instructions (BioRad). The synthesized cDNA served as the template for subsequent qPCR analysis. Specific primers for target DUB genes spotted on 96-well plate were used in the qPCR reactions. The qRT-PCR was run on a C1000 Touch Thermal Cycler (BioRad) and analyzed using CFX Maestro™ Software (BioRad). The relative expression levels of the DUB transcripts were calculated using the comparative Ct method as we did previously (78, 79) with normalization to housekeeping genes such as GAPDH, Rpl27, Hprt, and β-actin to account for variability in RNA input and reverse transcription efficiency.

### 4.6 Gentamicin protection assay

Cells were seeded at a concentration of 1.7 × 10^5 cells/mL and incubated for 48 hours prior to infection. On the second day, *Salmonella* Typhimurium UK-1 was prepared with 25 μg/mL chloramphenicol in LB Miller broth and incubated overnight at 37 °C with shaking at 250 rpm. On the day of infection, a subculture of S. Typhimurium UK-1 was prepared at an OD_600_ of 0.05 and grown for three hours at 37°C with shaking at 250 rpm. The macrophages were infected with the bacteria from the three-hour subculture in incomplete DMEM and incubated at 37 °C with 5% CO2 for one hour. Post-infection, the media was aspirated, cells were washed once with 1X DPBS, and incomplete DMEM supplemented with 100 μg/mL gentamicin was added. The cells were then incubated for an additional hour. Following this incubation, the media was aspirated, cells were washed once with 1X DPBS, and compounds diluted in incomplete DMEM with 25 μg/mL gentamicin were added. The cells were then incubated for the desired duration (2 to 24 hours).

### 4.7 Growth curve analysis

An overnight of Wild-type *Salmonella* Typhimurium UK-1 was prepared on the first day in 12 mLs of LB Miller and grown in a 37 °C shaking incubator set to 250 rpm. The next day, a subculture was prepared from the overnight culture at an optical absorbance of 0.05 similar to the subcultures prepared in the methodologies listed above. Enough subculture was prepared to fill a 96-well plate with 200 μL per well, and a concentration of 10 μM was prepared for each compound along with the appropriate controls – DMSO for the vehicle control, 100 μg/mL of ampicillin for the positive control, and an uninfected media-only control to serve as the blank for absorbance readings. The plate was then read using a Cytation3 (Agilent) set to 37 °C with a continuous shaking cycle, with OD_600_ readings taken at the start of the experiment and at every 30-minute interval over 24 hours. Absorbances were recorded using Excel (Microsoft) and analyzed relative to the vehicle control. The growth data was analyzed using R Studio (ver. 4.3.0). The analysis involved calculating the average and standard deviation of the replicates for each sample. The data was visualized using a bar plot with error bars indicating standard deviations. Samples with significant changes in growth were highlighted, and a dashed blue line was used to indicate 100% growth, serving as a reference point. The following R packages were used for the analysis: ggplot2, *dplyr*, and *tidyr*.

### 4.8 TNF-α ELISA

Cells were infected with wild-type *Salmonella* as described above, and media were collected 24 hpi and analyzed by the anti-TNF-α ELISA (Invitrogen). The data were analyzed using GraphPad Prism software. TNF-α concentrations were calculated based on a standard curve generated from known concentrations of TNF-α provided with the ELISA kit. Data were normalized to the "Vehicle Control" and pairwise t-tests were performed to compare TNF-α levels of each compound against the "Vehicle Control." The Benjamini-Hochberg correction was applied to control the false discovery rate. A bar graph was generated using the *ggplot2* package in R. Significance stars were added for compounds with adjusted *p*-values less than 0.05.

### 4.9 Western blot analysis

After following the gentamicin protection assay protocol listed above, the media we aspirated off the desired cells for Western blotting then washed twice with 1X PBS and left in 1X PBS. Using a cell scraper, cells were detached from the growing surface and collected in individually labeled sterile 1.5-mL Eppendorf tubes. Cells were spun down at 500 x g for 5 minutes to pellet the cells then PBS aspirated from the pellet without disturbing it. Pellets were resuspended in 120 μL of 1X radioimmunoprecipitation assay (RIPA) buffer to lyse the cells and incubate on ice or at 4 °C for 30 minutes with vortexing at every 5-minute interval. Next, the cells were placed in a sonicating water bath for 10 minutes to maximize protein yield. The lysed cell pellets were then spun at 15,000 x g for 10 minutes to pellet cell debris. The protein concentration was determined using a Bicinchoninic Acid (BCA) Protein Assay Kit (Thermo Scientific). Protein samples were separated using a 4 to 12% gradient SDS-polyacrylamide gel and transferred onto a polyvinylidene difluoride membrane (BioRad). The membrane was then blocked using 5% skim milk in Tris-buffered saline (TBS) with 0.1% Tween with proteins of interest being detected via immunodetection using appropriate antibodies and enhanced chemiluminescence. The antibodies used were specific for NF-κB (Cell Signaling, 8242T), phosphorylated NF-κB (p-NF-κB) (Cell Signaling, 3033S), p44/42 MAPK ERK1/2 (Cell Signaling, 4695T), p-p44/42 MAPK ERK1/2 (Cell Signaling, 4370T), TRAF3 (Cell Signaling, 4729T), USP25 (Santa Cruze Biotechnology, sc-398414), and β-actin (Santa Cruze Biotechnology, sc-47778). Secondary antibodies used for these Western blots were the goat anti-rabbit IgG (H+L) (Invitrogen, 31460), the goat anti-mouse IgG (H+L) (Invitrogen, 31430), and the mouse IgG1 BP-HRP (Santa Cruze Biotechnology, sc-525408) for use with the USP25 antibody, specifically.

### 4.10 QUANTI-Blue assay with RAW 264.7-Blue cells

RAW-Blue cells (InvivoGen) were used to perform a gentamicin protection assay following a protocol similar to the one described above. After the infection period and subsequent treatment with 100 µg/mL gentamicin, cells were incubated in 200 µL of testing media. At 4, 8, and 26 hours post-infection, as well as at a baseline time point prior to infection, 20 µL of supernatant was collected from the treated cells and mixed with 180 µL of prepared QUANTI-Blue solution, prepared according to the manufacturer’s specifications (InvivoGen). The supernatant/QUANTI-Blue mixture was incubated at 37 °C for 2 hours to allow color development and then analyzed for optical density at 620 and 655 nm using a Cytation3 plate reader.

For theLPS studies, RAW-Blue cells were incubated with either a vehicle control or AZ-1 in the presence or absence of 500 ng/mL LPS (Lipopolysaccharides from *Salmonella enterica* serotype Typhimurium | L6143, Millipore Sigma) in 200 µL of media. Supernatants were collected at 4, 8, and 26 hours post-inoculation, as well as at a baseline time point prior to inoculation. For each time point, 20 µL of supernatant was incubated with 180 µL of QUANTI-Blue solution for 16 hours at 37 °C before being read for optical density at 620 and 655 nm using the Cytation3 plate reader, as described above..

### 4.11. Cytotoxicity and viability assay

The cytotoxicity and viability of RAW 264.7 cells treated with Bortezomib (10 μM), AZ-1 (10 μM), or DMSO (vehicle control) after 24 hours and digitonin (positive control for complete lysis) after 15 minutes were evaluated using the Cytation 3 (Agilent) multifunctional reader. The MultiTox-Fluor Multiplex Cytotoxicity Assay was employed to determine the ratio of cytotoxicity to viable cells. Data are presented as mean ± SEM from three independent experiments and were analyzed using a one-way ANOVA followed by Tukey’s post-hoc test for multiple comparisons.

### 4.12. Murine experiment

#### Preparation of bacteria

An overnight culture in 5 ml LB Lennox plus 30 µg/mL Nalidixic Acid was made from frozen bacterial stock using a sterile pipette tip. After 16-18 hours of growth at 37°C with shaking at 220 rpm, a 20 mL sub-culture was made in fresh LB Lennox media from this overnight. This sub-culture was grown for 2-3 hours in the same condition as the overnight. After this time, the OD_600_ was measured and used to calculate the volume of culture necessary for the desired concentration of bacterial inoculum. *Salmonella* was spun at 10,000 x g for 10 minutes at 4°C to collect bacterial cells. Pellet was washed three times with 1X PBS before suspending in the required inoculum volume.

Stock AZ1 compound was prepared to a final concentration of 100 mM in DMSO. For a dose of 20 mg/kg, 7.1 µL of stock AZ1 was added to 100 µL of sterile 1X PBS. For the vehicle control dose, 7.1 µL of DMSO was added to 100 µL of sterile 1X PBS.

C57BL/6J (000664) mice were acquired from University of Florida Animal Care Services (ACS) Rodent Models Breeding Core. Female, age-matched mice of 8 weeks were used for the experiment in this study. Control mice and experimental mice were randomly selected for treatments. All mice were housed in a specific pathogen-free (SPF) animal facility at University of Florida’s Department of Veterinary Medicine with a 12-hr dark/12-hr light cycle and fed standard chow and water. Upon arrival, mice were acclimated to the housing conditions before experimental treatments began. Baseline weights were collected prior to any experimental treatment. For identification within cages, ear punches were utilized to properly differentiate and track each mouse. On Day 2 post-infection, mice were provided a soft chow to assist in extending survival. Mice were administered a vehicle control (DMSO in PBS) or AZ1 (20 mg/kg body weight) in 100 µL by oral gavage daily. Death of a mouse or a 20% reduction in body weight signified the endpoint of the study.

24 hr prior to inoculation of *Salmonella* Typhimurium 12023 Nalidixic Acid (NA) Resistance, 8-week-old mice were administered 100 µL of freshly prepared 200 mg/mL streptomycin sulfate in sterile water via oral gavage. This antibiotic solution was allowed to establish in the gut of the mice for 24 hours to suppress gram-positive bacteria resides in the gut microbiome. The next day, each mouse was administered a dose of 1 x 10^8^ CFU of bacteria in 50 µL via oral gavage. 10 minutes prior to inoculation of mouse with bacteria, 0.3 M Sodium Bicarbonate in 50 µL was administered via oral gavage to neutralize the stomach acid and protect viability of the inoculum. *Salmonella* was allowed to establish overnight before the first dose of AZ1 treatment or vehicle control was administered using oral gavage. Mice were administered AZ1 or vehicle control daily until the endpoint of the study.

Additionally, fecal samples were taken on Day 2 post-infection for assessment of bacterial load and inflammatory response, respectively. For fecal analysis of salmonellae, sterile 1.5 mL microcentrifuge tubes were weighed to obtain each tube’s weight. Mice from different experimental groups were separately placed in a sterile pup cup. Mice were left in pup cups until droppings were able to be collected. Using a sterile 10 µL pipette tip, fecal dropping were collected and transferred into the sterile, weighed 1.5 mL microcentrifuge tubes. The weight of the tube plus fecal sample was calculated to determine the weight of each fecal sample. Following this, 1 mL of sterile 1X PBS was added to each feces-containing tube and subjected to homogenization on a vortex at setting 7 for 15 minutes. Serial dilutions were performed of the fecal slurry using 30 µL of samples into 270 µL of sterile 1X PBS. 100 µL of the dilutions 10^-1^, 10^-4^, and 10^-5^ were plated in triplicate for each mouse on LB Lennox agar containing 30 µg/mL Nalidixic Acid. Plates were incubated overnight at 37°C. The number of CFUs was recorded and used to determine the CFU/g for all mice.

## 4. Acknowledgements

We would like to thank Dr. Anna Aulicino and Dr. Alison Simmons (University of Oxford, UK), for their valuable discussions and comments. We would like to thank Dr. Manuela Manuela Raffatellu (UC San Diego) for *Salmonella* Typhimurium 12023 with Nalidixic Acid resistance. This study was funded by the Department of Defense grant (W81XWH-20-1-0176) and U.S. National Institute of Allergy and Infectious Diseases (NIAID) via R01 AI158749-03 (M.J.F.).

